# Case Report: Aggressive Tibial Pseudarthrosis as Primary Symptom in Infant with Neurofibromatosis

**DOI:** 10.1101/066316

**Authors:** Mary Alice Reid, Madeline Zupan, Nicole Sevison, Barbara Calhoun, Kasturi Haldar

**Affiliations:** Beacon Health System, Bristol Street Pediatrics, Goshen, Indiana; Boler-Parseghian Center for Rare and Neglected Diseases, Dept. Biological Sciences, University of Notre Dame Notre Dame, Indiana; Memorial Hospital, South Bend, IN

**Author notes:** **Address correspondence to**: Dr. Kasturi Haldar, PhD, Boler-Parseghian Center for Rare and Neglected Diseases, Dept. Biological Sciences, 107 Galvin Life Sciences Building, Notre Dame, Indiana, [ ].

## Abstract

Neurofibromatosis (NF1) is a rare genetic neurologic disorder with over 30 distinct clinical manifestations, the top 3 of which include café-au-lait spotting, benign tumors and abnormal freckling. Pseudarthrosis (PA), also known as a “false joint,” is a rare subset of NF1 symptomology, characterized by bone fractures and nonunion caused by severe bowing of long bones. To date, it is invariably reported as secondary to NF1, commonly at 24 months of age. Here we describe a 4-month old infant who presented with PA as primary symptom, and in absence of an NF1 first-degree relative. Initial manifestation was guarding of the leg and increased irritability upon palpation of the knee, subsequent to light playful jostling. Physician examination revealed gross anterolateral bowing of the left leg. Radiography confirmed tibia-fibula bowing and pathologic transverse fracture at tibia diaphysis, characteristic of PA. Café-au-lait spots developed at 6 months subsequent to PA, but with number and size well below the National Institutes of Health criteria for NF1 diagnosis. At 14 months, exome sequencing established definitive NF1 diagnosis. Treatment involved PA takedown surgery. Although healing was seen after 2 months, complications emerged by 6 months. This case suggests that for primary PA without clear etiology, first-contact and consulting physicians should pay careful attention and be vigilant to timing of clinical onset and severity. Early, severe primary PA warrants accelerated NF1 exome sequencing, suggesting expansion of existing federal guidelines may be necessary to improve detection and prognosis of this rare, debilitating but readily managed condition.

**Financial Disclosure:** The authors have no financial relationships relevant to this article to disclose

**Conflict of Interest:** The authors have no conflicts of interest to disclose

**Clinical Trial Registration:** NA

**Abbreviations**
(NF1): Neurofibromatosis 1
(PA): Pseudarthrosis
(NIH): National Institutes of Health

## Contributors’ Statements

Kasturi Haldar conceived and designed the overall study, undertook data interpretation and analysis and provided intellectual framework and content in writing and revising all drafts of the manuscript. Mary Alice Reid conceived and designed the study, provided patient clinical data, its analysis and interpretation and critically revised drafts for intellectual content. Madeline Zupan undertook data analysis and provided intellectual content in writing and revising all drafts of the manuscript. Barbara Calhoun conceived and designed the study and provided intellectual content in revising drafts of the manuscript. Nicole Sevison provided consultation and patient clinical data, undertook data analysis, and provided intellectual content in revising the manuscript. All authors approved the final manuscript as submitted and agree to be accountable for all aspects of the work.

## Introduction

NF1 (OMIM #162200) is a rare autosomal dominant genetic condition that targets the nervous system resulting in neurofibromas, typically benign tumors that affect the brain, spinal cord, nerves, and skin. Disease incidence estimations range from 1 in 2,500 to 4,000 births and affect males and females in equal frequencies. Individuals may genetically inherit the disorder, but approximately 50% of mutations occur *de novo* in the *NF1* gene encoding neurofibromin tumor suppressor protein on chromosome 17q11.2^1^. The mutation rate at this locus is among the highest rate known for any gene in humans.

This degenerative disease is characterized by a wide spectrum of clinical manifestations such as anomalous skin hyperpigmentations (café-au-lait spotting), neurofibromas, bone growth abnormalities, learning deficits, and developmental delays. The clinical symptoms are diverse in phenotypic manifestation as well as symptomatic severity. These factors present challenges to the diagnosis of NF1, which is contingent upon guidelines enumerated by the National Institutes of Health (NIH). The NIH specifies that NF1 should be *suspected* if one of the following criteria are present and *diagnosed* if any *two* of the following seven diagnostic criteria are met: six or more café-au-lait spots over 15 mm in diameter in post-pubertal individuals or 5 mm in pre-pubertal individuals, two or more neurofibromas (of any type) or one plexiform neurofibroma, axillary or inguinal freckling, optic gliomas, two or more iris hamartomas (Lisch nodules), osseous lesions such as sphenoid dysplasia or PA, or an NF1 diagnosed first-degree relative^2^. A multistep mutation detection cDNA sequence analysis is available that definitively detects greater than 95% of NF1 pathogenic variants^1,3-5^.

PA is a distinct orthopedic manifestation primarily seen as secondary to NF1. It also has diverse clinical presentations, ranging from anterolateral tibial bowing to disjointed or nonunion fractures. This subset is a particularly rare manifestation, as only 2% of individuals with NF1 develop PA and/or bowing of the long bones^6,7^ symptomatically manifest within the first few years of life^8^. Here, we present the first reported case of PA as *primary* manifestation of NF1 in an infant female at only four months of age.

## Case Report

A pediatrician evaluated a female four-month old after patient’s mother noted the infant was irritable and guarding left leg against abdomen. The baby unexpectedly expressed increased pain with diaper changes and palpation of the knee area. The baby was born via normal (38 weeks) gestation and delivery after an uneventful pregnancy. Apgar scores were 8 (1 minute) and 9 (5 minutes). Birth weight was 5 pounds 7 oz., height 19”, and head circumference 12.99’’. Infant was noted to have jaundice that resolved after one week (Figure 1). Baby breastfed regularly, but four-month visit indicated patient maintained height at fifth percentile and weight at third percentile. Patient met all other developmental milestones including babbling, recognizing parent’s voice, lifting head when prone, and controlling head while sitting at four-month visit.

**Figure 1.**
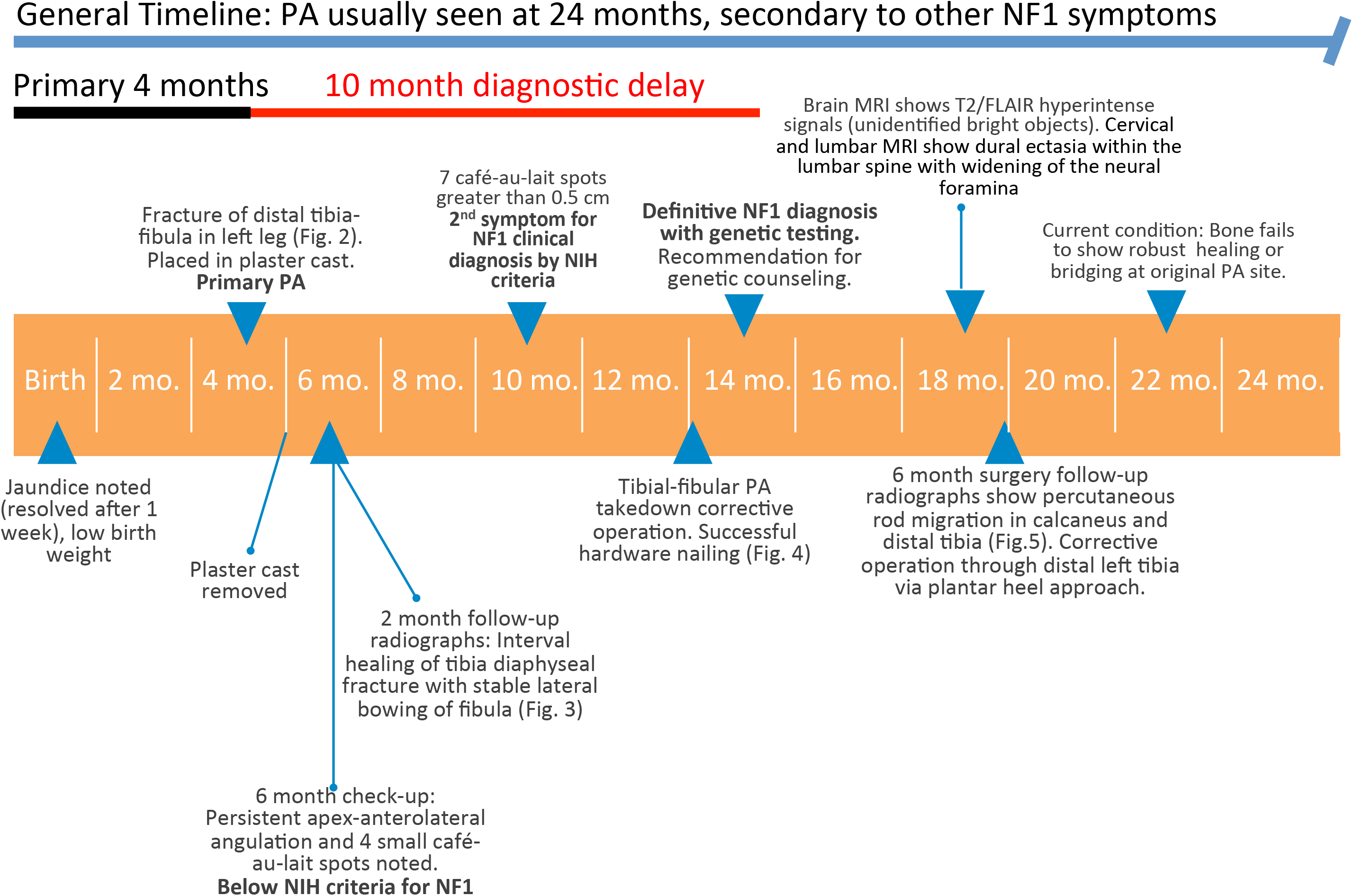
Diagnostic timeline of the case report. Patient presented primary symptom at 4 months but did not establish definite diagnosis until 14 months only with genetic testing. Accelerated NF1 exome sequencing could lead to better diagnosis of ulterior problems and improved management of patient outcomes.

Fracture occurred at four months of age (Figure 1) when patient’s father was gently bouncing child on knee and noticed abrupt change in behavior as patient began to scream and guard the left leg. Patient was still guarding lower extremity when assessed the next day by pediatrician. Radiological images of distal left tibia-fibula were suspicious for neoplastic process secondary to tibia fracture and bowing deformity of tibia and fibula (Figure 2). Patient was further evaluated with a CT scan of leg two days later that revealed severe anterolateral bowing of the left tibia and fibula (not shown, summarized in Figure 1). Findings revealed sequelae of isolated congenital PA of the tibia. Patient was placed in long-leg plaster cast for four weeks. Two-month follow-up radiology showed interval healing of the tibial diaphyseal fracture and slight lateral angulation at the fracture apex with stable lateral bowing of the fibula (Figure 3).

**Figure 2.**
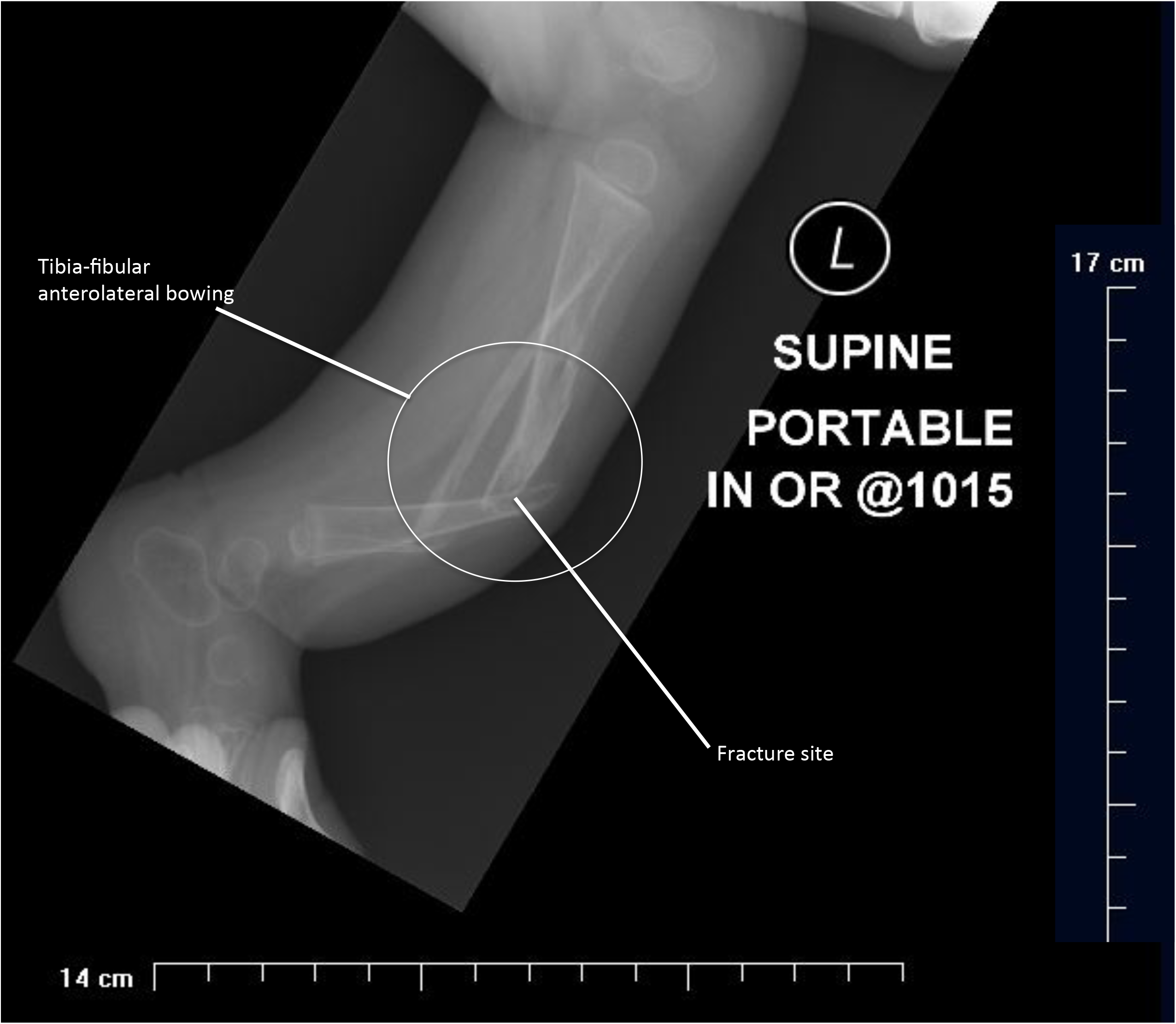
Post-fracture radiographs showing anterolateral bowing of the left tibia-fibula in 4-month old pseudarthrotic infant with neurofibromatosis 1. Anterior view of left tibia and fibula with pathologic transverse nondisplaced fracture and lateral anterior bowing at the distal one-third of the diaphysis.

**Figure 3.**
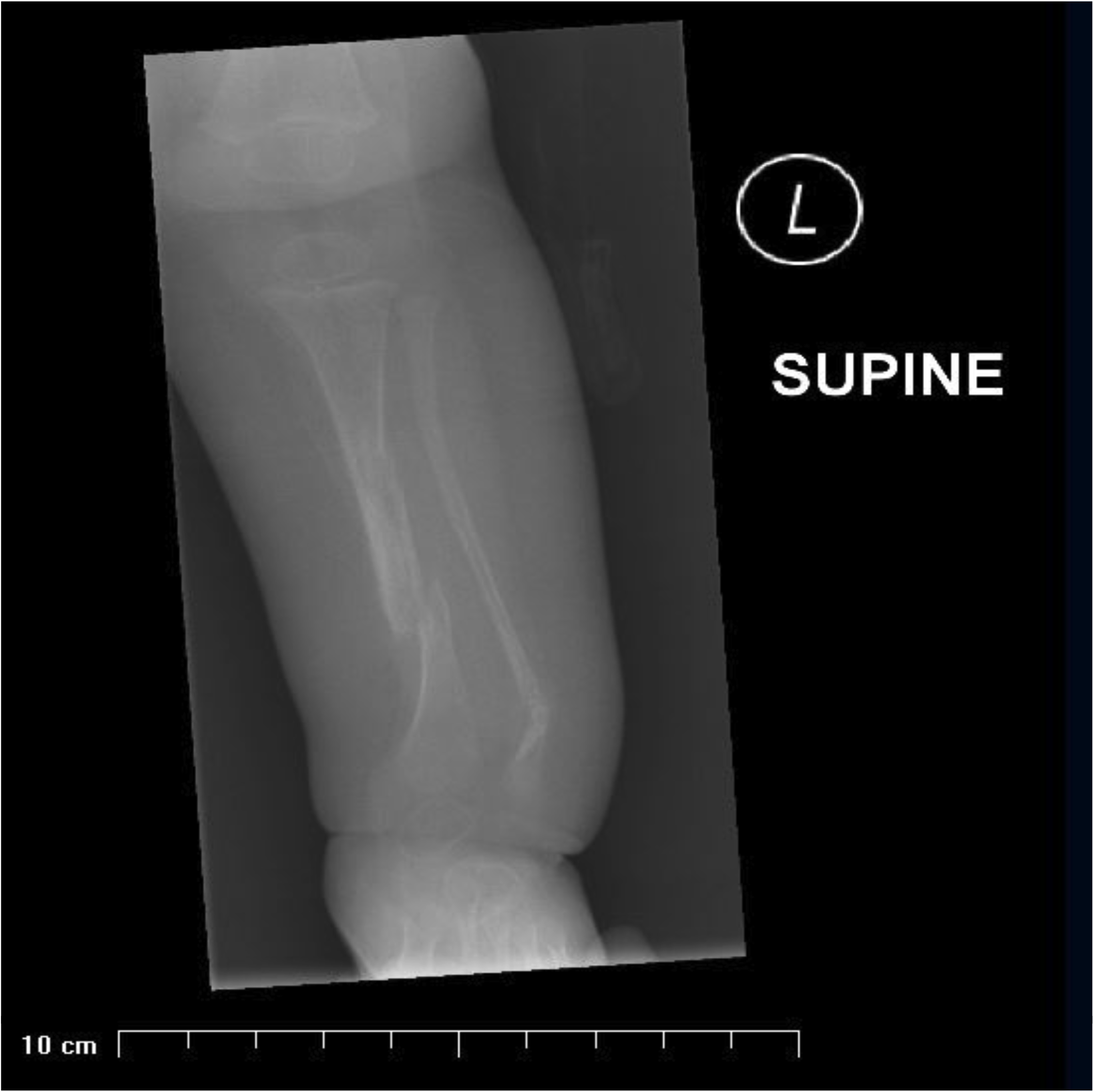
Pre-operative radiograph showing anterolateral bowing of the left tibia-fibula. Two-month fracture follow-up radiology showing medial view of left tibia and fibula indicates interval healing of the tibial diaphyseal fracture and slight lateral angulation at the fracture apex.

At the six-month check-up, patient exam revealed persistent lower left leg apex-anterolateral angulation with a bony mass at the PA location and four café-au-lait spots (Figure 1). Spot count was still below the number of six café-au-lait spots of 5 mm acceptable as a second diagnostic criterion for suspecting NF1, as per NIH. By ten months of age, café au-lait-spot count increased to seven, all with measurement greater than 5 mm, suggesting that six months after the primary presentation of PA, patient met NIH criteria for suspecting NF1 (Figure 1).

At thirteen months, patient underwent first corrective operation with takedown of left tibial-fibular congenital pseudarthrosis, iliac crest bone graft, and intramedullary rod nailing (Figure 4). A one and one-half hip spica cast was applied and patient followed up monthly with orthopedist. Confirmation of NF1 diagnosis occurred one month later with a molecular genetic test yielding a heterozygous truncating mutation at 17q11.2, exon 40 on the *NF1* gene (summarized in Figure 1). No other mutational alteration was detected with *NF1* mutation analysis.

**Figure 4.**
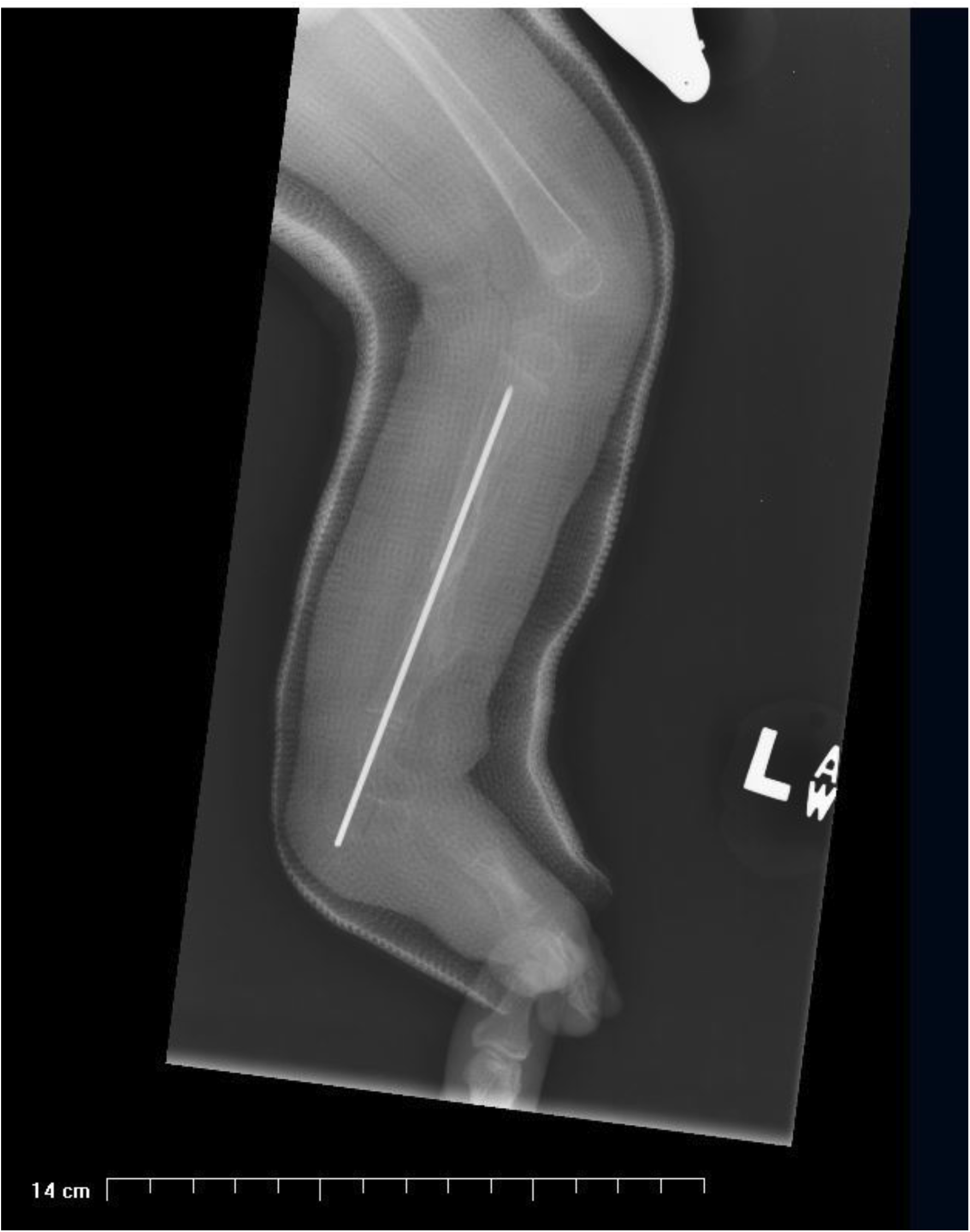
Post-operative radiograph of intramedullary rod in left tibia. Surgical intervention of left tibial-fibular congenital pseudarthrosis takedown, iliac crest bone graft, and intramedullary nailing. A 1-1/2 hip spica cast was applied and radiographs indicate a successful operation.

At patient age of 18 months, brain MRI testing revealed a T2/FLAIR hyperintense signals (unidentified bright objects) along the medial right and left cerebellar hemispheres along the posterior lateral aspect of the fourth ventricle (not shown, summarized in Figure 1). Cervical and lumbar MRI revealed dural ectasia within the lumbar spine with widening of the neural foramina bilaterally throughout the lumbar spine and minimal posterior vertebral body scalloping in the lumbar vertebral bodies (not shown, summarized in Figure 1). These symptoms are all characteristic of NF1.

At patient age of nineteen months, six-month surgery follow-up radiographs showed significant migration of the inserted percutaneous rod hardware. The most distal aspect of the rod in the inferior tibia migrated to the plantar skin overlying the calcaneus (Figure 5). Moderate lucency about the calcaneus was noted, indicating significant rod motion. The area of PA in the mid-diaphysis of the tibia did not show any significant regeneration of bone in the graft site and the position of the bone structures remained unchanged. A corrective operation was immediately undertaken through distal left tibia via a plantar heel approach (Figure 1).

**Figure 5.**
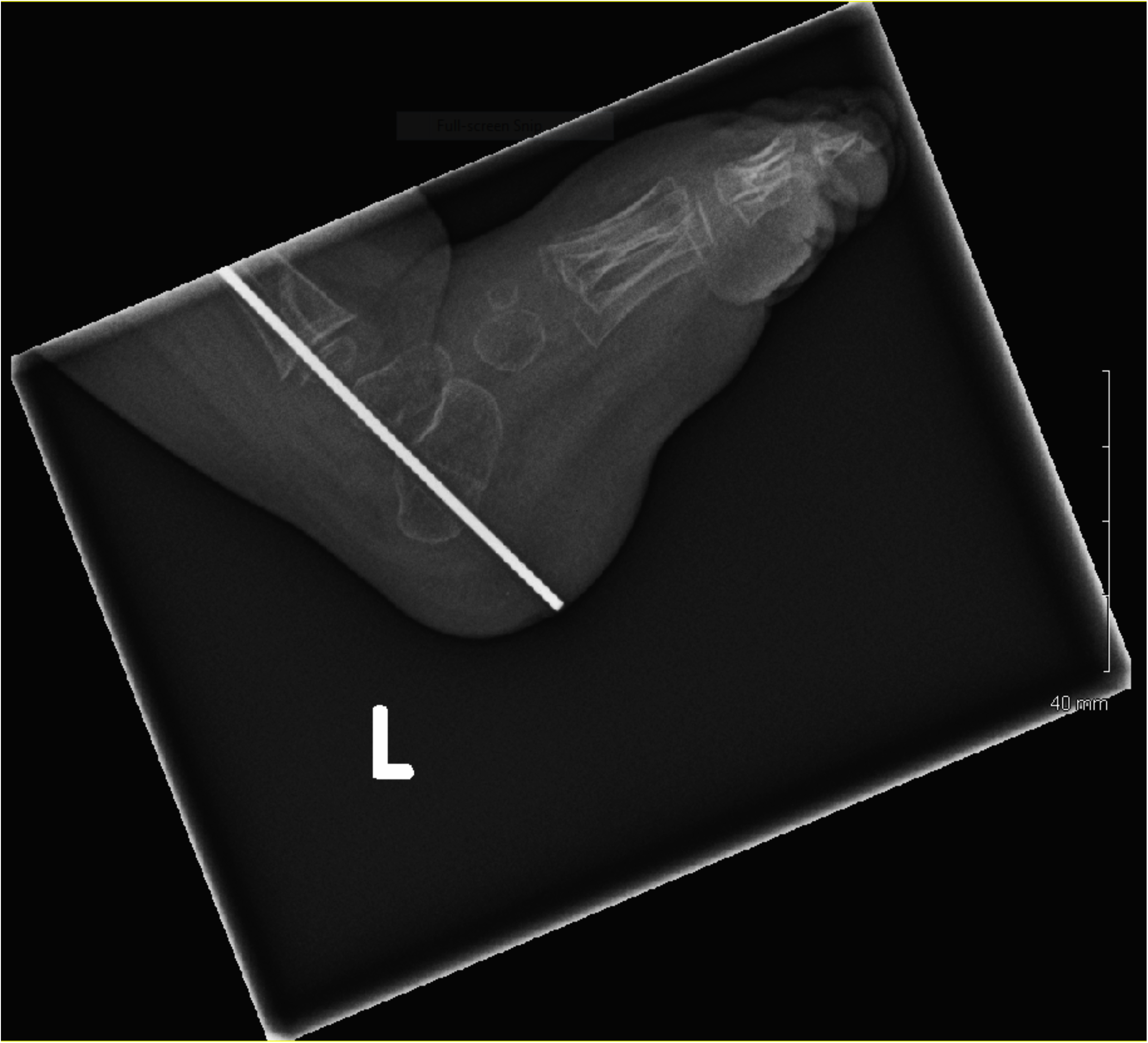
Intramedullary hardware rod migration. Six month post-operative radiographs shows the distal-most aspect of the rod is at the plantar skin overlying the calcaneus indicating significant hardware migration post-surgery at patient age of 19 months. Patient underwent immediate corrective operation. Rod length shortened and re-fixated via a plantar heel approach.

At present condition, twenty-two month pediatric and radiologic evaluation shows stable but slow healing of fracture and progressive straightening of leg. Radiographs show improved solidified osteogenesis at the proximal tibia, however bone fails to show robust healing or bridging at the original PA site (not shown, summarized in Figure 1). Patient shows significant gross physical development delays (<5^th^ percentile weight, <5^th^ percentile length, <28^th^ percentile head circumference), failure to thrive, frontal bossing, elongated digits, T2/FLAIR hyper-intense cranial signals, and seven café-au-lait spots greater than 5 mm in size. NF1 patients often also have ophthalmologic manifestations but currently patient reports negative for these NF1 clinical symptoms. Patient family history is negative for genetic bony dysplasias and neurologic disorders. Treatment regimen is full-time use of a knee-ankle-foot orthotic and daily ultrasound bone stimulation.

## Discussion

NF1 patient treatment options are limited as no definitive FDA-approved cure or therapy exists. Symptom management is typically conducted by specialist referral for ocular, neurological, or osteological abnormalities. PA pathogenesis is a result of fibrous hamartoma causing diminished mechanical bone strength and osteogenesis. High levels of cellular fibrovascular tissue grow into the bony cortex region, negating normal bone modeling^9^. At the medullary bone level, reactive changes lead to the deposition of excess trabecular bone causing medullary sclerosis^9^. Both osteoblasts and osteoclasts function abnormally in people with NF1^10,11^ and impaired vascularization diminishes any further osteogenesis^9^. Treatment is difficult because once bone fracture occurs, it is challenging to achieve spontaneous regrowth and autonomous healing fails. Despite corrective operations, re-fracture and growth disturbances are frequent, as observed in this case^8^. Recombinant human bone morphogenetic protein (rh-BMP) is currently under investigation as a potential therapeutic option for pseudarthrosis and other long bone fracture candidates^9,12^.

This case is of particular medical interest due to the unusual phenotype of the neurofibromatosis. The disorder manifested itself in our patient predominantly through orthopedic – rather than neurologic – symptoms. Furthermore, this case is unique in that initial fracture occurred at four months of age, indicating a particularly aggressive congenital disorder despite absence of prior family history of either PA or NF1. Our patient was a small, light infant not bearing full weight on her long bones. Due to the severely dysplastic and bowed nature of her tibia and fibula, fracture occurred with minimal force (presumably the downward bounce on the bowed leg). Such minimal activity typically does not yield fractures especially in normal infants. This case illustrates one extreme end of the spectrum for bone dysplasia severity.

Since in prior reported cases of NF1, PA was invariably secondary to NF, our patient did not present enough early symptoms to receive NF1 diagnosis at 4 months when she first manifested PA. Up to one-half of children who are first seen by orthopedists for anterolateral bowing of the tibia will not have a diagnosis of neurofibromatosis^13^. Although the NIH criteria for suspecting NF1 provide high value, there is a need for further improvement, as up to 35% of children with NF1 remain undiagnosed by the age of five^14,15^. For example, one of the primary diagnosis criterions stipulates multiple café-au-lait spots yet 10% of the general healthy population has one or two café-au-lait spots^6^. Due to these overlapping symptoms, NF1 diagnosis becomes muddled and may only be definitively established with exome sequencing. It is thus our recommendation that any suspected case of NF1 - due to particular severe, early, or primary PA - may benefit from accelerated NF1 exome sequencing leading to better diagnosis of ulterior problems and improved management of patient outcomes. Therefore expansion of existing NIH guidelines is warranted to accommodate such aggressive cases to improve pre-symptomatic detection of treatable manifestations, reduce risk of fracture prior to breakage and to improve ultimate prognosis of this rare condition.

